# The colonial legacy of herbaria

**DOI:** 10.1101/2021.10.27.466174

**Authors:** Daniel S. Park, Xiao Feng, Shinobu Akiyama, Marlina Ardiyani, Neida Avendaño, Zoltan Barina, Blandine Bärtschi, Manuel Belgrano, Julio Betancur, Roxali Bijmoer, Ann Bogaerts, Asunción Cano, Jirí Danihelka, Arti Garg, David E. Giblin, Rajib Gogoi, Alessia Guggisberg, Marko Hyvärinen, Shelley A. James, Ramagwai J. Sebola, Tomoyuki Katagiri, Jonathan A. Kennedy, Tojibaev Sh. Komil, Byoungyoon Lee, Serena M.L. Lee, Donatella Magri, Rossella Marcucci, Siro Masinde, Denis Melnikov, Patrik Mráz, Wieslaw Mulenko, Paul Musili, Geoffrey Mwachala, Burrell E. Nelson, Christine Niezgoda, Carla Novoa Sepúlveda, Sylvia Orli, Alan Paton, Serge Payette, Kent D. Perkins, Maria Jimena Ponce, Heimo Rainer, L. Rasingam, Himmah Rustiami, Natalia M. Shiyan, Charlotte S. Bjorå, James Solomon, Fred Stauffer, Alex Sumadijaya, Mélanie Thiébaut, Barbara M. Thiers, Hiromi Tsubota, Alison Vaughan, Risto Virtanen, Timothy J. S. Whitfeld, Dianxiang Zhang, Fernando O. Zuloaga, Charles C. Davis

**Affiliations:** Department of Biological Sciences, Purdue University, West Lafayette, IN, USA; Purdue Center for Plant Biology, Purdue University, West Lafayette, IN, USA; Department of Geography, Florida State University, Tallahassee, FL, USA; Department of Botany, National Museum of Nature and Science, Tsukuba, Ibaraki, Japan; Herbarium Bogoriense, Research Center for Biology, National Research and Innovation Agency (BRIN), Cibinong 16911, West Java, Indonesia; Instituto Experimental Jardin Botánico “Dr. Tobías Lasser”, Avenida Salvador Allende, Ciudad Universitaria, Caracas, Venezuela; Universidad Central de Venezuela, Caracas, Venezuela; Hungarian Natural History Museum, Budapest, Hungary; CeReSE - Université Claude Bernard Lyon 1, Lyon, France; Instituto de Botánica Darwinion, Acassuso, Provincia de Buenos Aires, Argentina; Universidad Nacional de Colombia, Bogotá, Colombia; Naturalis Biodiversity Center, Botany Section, Leiden, Netherlands; Meise Botanic Garden, Meise, Belgium; Herbario San Marcos, Museo de Historia Natural, Universidad Nacional Mayor de San Marcos, Lima, Peru; Department of Botany and Zoology, Faculty of Science, Masaryk University, Brno, Czech Republic; Institute of Botany of the Czech Academy of Sciences, Průhonice, Czech Republic; Botanical Survey of India, Central Regional Centre, Allahabad, India; University of Washington Herbarium, Burke Museu, Seattle, WA, USA; Botanical Survey of India, Sikkim Himalaya Regional Centre, Sikkim, India; Institute of Integrative Biology, ETH Zurich, Zurich, Switzerland; Botany Unit, Finnish Museum of Natural History, University of Helsinki, Helsinki, Finland; Department of Biodiversity, Conservation and Attractions, Western Australian Herbarium, Kensington, WA, Australia; South African National Biodiversity Institute, Pretoria, South Africa; School of Animal, Plant and Environmental Sciences, University of the Witwatersrand, Johannesburg, South Africa; Hattori Botanical Laboratory, Nichinan, Miyazaki, Japan; Harvard University Herbaria, Cambridge, MA, USA; Institute of Botany, Uzbekistan Academy of Sciences, Tashkent, Uzbekistan; National Institute of Biological Resources, Incheon, South Korea; National Parks Board, Singapore Botanic Gardens, Singapore; Department of Environmental Biology, Sapienza University of Rome, Rome, Italy; Herbarium Patavinum, University of Padua, Padua, Venetia, Italy; East African Herbarium, National Museums of Kenya, Nairobi, Kenya; Komarov Botanical Institute, Russian Academy of Sciences, Saint Petersburg, Russian Federation; Herbarium collections & Department of Botany, Faculty of Science, Charles University, Prague, Czechia; Institute of Biological Sciences, Maria Curie-Skłodowska University, Lublin, Poland; Rocky Mountain Herbarium, University of Wyoming, Laramie, WY, USA; Field Museum, Chicago, IL, USA; Staatliche Naturwissenschaftliche Sammlungen Bayerns, Botanische Staatssammlung München, Germany; Department of Botany, National Museum of Natural History, Smithsonian Institution, Washington, DC, USA; Royal Botanic Gardens, Kew, Surrey, United Kingdom; Herbier Louis-Marie, Université Laval, Québec, Canada; University of Florida Herbarium, Florida Museum, Gainesville, FL, USA; Instituto Multidisciplinario de Biología Vegetal, (UNC-CONICET) Córdoba, Argentina; Naturhistorisches Museum Wien, Wien, Austria; Department of Botany and Biodiversity Research, University of Vienna, Vienna, Austria; Botanical Survey of India, Deccan Regional Centre, Hyderabad, India; National Herbarium of Ukraine, M.G. Kholodny Institute of Botany, National Academy of Sciences of Ukraine, Kyiv, Ukraine; Natural History Museum, University of Oslo, Oslo, Norway; Missouri Botanical Garden, St. Louis, MO, USA; Conservatory and Botanic Gardens of Geneva, Geneva, Switzerland; Department of Plant Sciences, University of Oxford, Oxford, United Kingdom; The New York Botanical Garden, Bronx, NY, USA; Graduate School of Integrated Sciences for Life, Hiroshima University, Higashi-Hiroshima, Hiroshima, Japan; Miyajima Natural Botanical Garden, Hiroshima University, Hatsukaichi, Hiroshima, Japan; Royal Botanic Gardens Victoria, Melbourne, Australia; Ecology and Genetics Research Unit, University of Oulu, Oulu, Finland; University of Oulu Botanical Museum, Oulu, Finland; Bell Museum, University of Minnesota, St Paul, MN, USA; South China Botanical Garden Herbarium, Chinese Academy of Sciences, Guangzhou, China; Department of Organismic and Evolutionary Biology, Harvard University Herbaria, Harvard University, Cambridge, MA, USA

**Keywords:** biodiversity, colonialism, digitization, GBIF, herbarium

## Abstract

Herbarium collections shape our understanding of the world’s flora and are crucial for addressing global change and biodiversity conservation. The formation of such natural history collections, however, are not free from sociopolitical issues of immediate relevance. Despite increasing efforts addressing issues of representation and colonialism in natural history collections, herbaria have received comparatively less attention. While it has been noted that the majority of plant specimens are housed in the global North, the extent of this disparity has not been rigorously quantified to date. Here, by analyzing over 85 million specimen records and surveying herbaria across the globe, we assess the colonial legacy of botanical collections and how we may move towards a more inclusive future. We demonstrate that colonial exploitation has contributed to an inverse relationship between where plant biodiversity exists in nature and where it is housed in herbaria. Such disparities persist in herbaria across physical and digital realms despite overt colonialism having ended over half a century ago, suggesting ongoing digitization and decolonization efforts have yet to alleviate colonial-era discrepancies. We emphasize the need for acknowledging the inconvenient history of herbarium collections and the implementation of a more equitable, global paradigm for their collection, curation, and use.

## Main

The nearly 400 million specimens residing in the world’s herbaria form the basis of the scientific understanding of our planet’s flora and are a centerpiece of botanical research and discovery ^1^. Since the 16th century, scientists including Linnaeus and Darwin have collected herbarium specimens principally to describe species and circumscribe taxonomic classifications. The past decade has seen a resurgence in herbarium collections research, which is driven in part by massive digitization efforts ^2–4^. In particular, with advances in high-throughput methods and image analyses, herbarium specimens are increasingly being used in innovative ways ^5,6^ beyond their original intended purpose, including research pertaining to global change ^7–9^. For example, herbarium specimens have been used to uncover the effects of climate change on plant phenology ^10^, ecophysiology ^11^, and herbivory ^7^; as barometers for pollution ^12^ and eutrophication trends ^13^; and to reconstruct the origin and spread of invasive species ^14,15^.

However, these collections are not free from the many sociopolitical issues that define our modern era. Despite increased efforts by natural history museums and other cultural institutions to address their legacy of colonialism and representation, such efforts have largely been focused on human- and animal-related collections and public exhibits ^16,17^. In contrast, herbaria have received comparatively less attention, sidelined by their lower visibility; few herbaria offer public displays and plant awareness is generally lacking ^18^. Nonetheless, botanists have contributed significantly to the colonial expansion of imperial powers through active participation in the overseas collection of plants and their scientific and economic development ^19^. For instance, Hans Sloane (1660–1753), often credited as the inventor of chocolate milk, collected hundreds of plant specimens from overseas colonies via the slave trade, including those of cacao ^20^. In Australia and other colonies, much of the early exploration of the continent by colonialists (during which many botanical specimens were collected) was done with the assistance of Indigenous peoples who acted as guides during expeditions – often under duress – or were forced to disclose their scientific knowledge of plants and place ^21–23^. Though not specifically quantified, it has thus been noted that herbaria in the global north hold many of the voucher specimens and associated data from equatorial and southern hemisphere nations (i.e., the global south), owing to colonial-era explorations ^24,25^.

To address the past and present appropriation of plant diversity and to open a dialogue to help move us towards a more inclusive future, we must first understand the extent of disparity in herbarium collections across the globe – specifically, a more robust history of where they were collected and where they currently reside. Here, we, scientists and curators from herbaria across 40 countries from every continent, examine the colonial legacy of botanical collections by assessing the geopolitical distribution of herbarium collections and digitization efforts. Analyzing over 85 million plant specimen records from the Global Biodiversity Information Facility (GBIF, April 23, 2021; https://doi.org/10.15468/dl.nt5wkx), the largest repository of biodiversity data, and surveying herbarium staff an curators across the world, we provide an in-depth view of the disparity present in herbarium collections and discuss the future of this colonial legacy and how its effects can be mitigated (see Supplementary Information for discussions about methods and assumptions).

### The imprint of colonialism in online herbarium collections

Collection trends across the last four centuries strongly bear the imprint of colonialism, especially among European institutions. These trends can be readily observed in the plant specimen records hosted on GBIF, which represent a subset (∼25%) of global herbarium collections (Fig. 1A). Large numbers of plant specimens collected across the globe are currently housed in European countries, and to a lesser extent, the United States. Indeed, the currently widely adopted taxonomy of life originated from European scholars, most prominently Linnaeus and his acolytes, who were responsible for the relocation of massive numbers of plant collections from across the globe into European institutions and their associated systems of knowledge. This trend was further fueled by the desire by imperial powers to exploit the biological resources of colonies abroad, a legacy of which persists to this day, for instance in the pursuit of medicinal plants in tropical regions in search of profitable remedies for predominantly First World ailments such as cancer or obesity ^26^.

**Figure 1.**
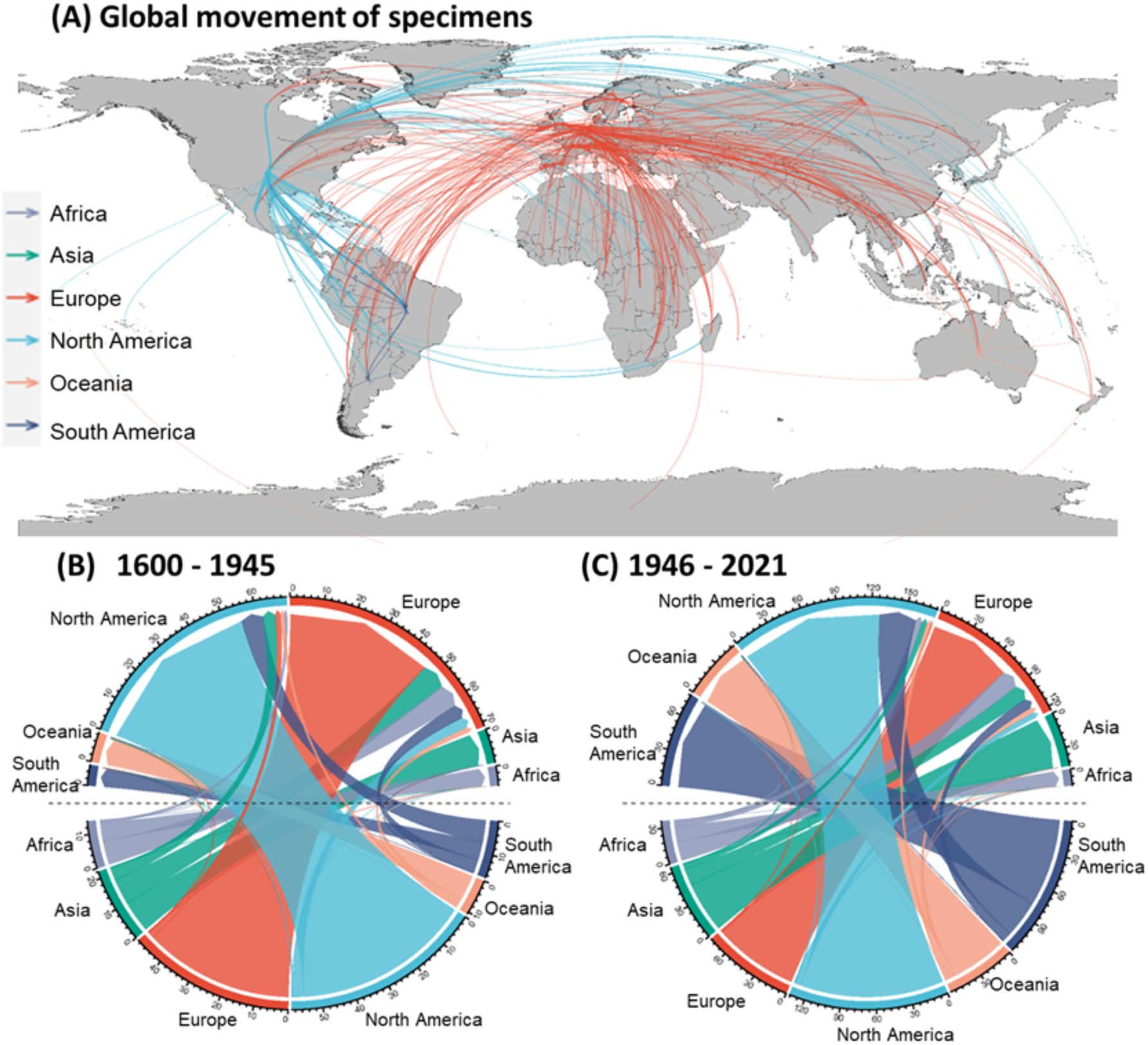
The past movement of plant specimens across the globe based upon records from GBIF. The world map depicts the top 10 percentile of intercontinental connections between countries where specimens have been collected and where they are currently housed regardless of collection date (A). The widths of the arrows are proportionate to the number of specimens dislocated and are colored by destination continent. Collections that remained in the country of collection are not depicted. The lower panels illustrate the intercontinental flow of specimens before (B) and after (C) the end of overt colonialism post World War II (late-1945). Arrows are colored by the continent of origin. Numbers on the outer ring indicate specimen numbers collected from (lower half) or stored in (upper half) each continent, and are in multiples of 100,000. Colors on the outer ring represent different continents.

The impact of this collecting legacy persists in the trends and patterns of more recent collecting activities. Despite the era of overt colonialism drawing to an end after the Second World War, the historical trend of specimen movement from Africa, Asia, and South America to Europe and North America has largely remained constant (Fig. 1B, C), especially among countries that have historical connections ^27^. In fact, the proportion of specimens collected from other continents has increased in Europe and North America over time. In particular, the United States emerged as the largest collector of overseas specimens after the Second World War, acquiring massive collections from countries such as Brazil and Madagascar. Notably, the proportion of specimens collected and housed in South America greatly increased during this period, while collection activity in Africa remained largely driven by European and North American countries, with the possible exception of South Africa. These trends are largely consistent when limiting our sample to records with more complete information (e.g., geographic coordinates; Supplementary Fig. 1) or to collection activity in the 21^st^ century (Supplementary Fig 2). However, we note that there are other factors, such as the degree of economic development, regional policies, political stability, and scientific interest that have likely influenced these patterns as well ^28^. Also, though difficult to estimate, a portion of the specimens that have been dislocated likely have duplicates deposited at local institutions, especially among more recent collections that may yet remain undigitized and thus less discoverable. Among the specimen data we examined from GBIF, only 2.8% were of the same species collected at the exact same place and date and stored in different institutions.

Our analysis clearly demonstrates that colonial exploitation has contributed to a striking inverse relationship between where plant biodiversity exists in nature versus where it is housed in herbaria. In general, biodiversity is distributed along a latitudinal gradient, with most of the world’s plant diversity located in the tropics ^30^. However, when we examine the number of species collected in a given country – which reflects species richness – relative to the number of species with specimens housed in the same country, striking disparities emerge (Fig. 2A). Specifically, most of the world’s flora is stored in temperate regions in a reverse-latitudinal gradient. In particular, herbaria in the United States and several nations in western and central Europe house over twice the number of species that occur in these nations, demonstrating the international appropriation of large amounts of plant diversity (Fig. 2A). In contrast, much of Africa and Asia house fewer species than are collected there, because North American and European herbaria currently house much of the specimens and associated data from these regions owing, in no small degree, to their colonial past. Indeed, nations from these two areas simultaneously i) house a disproportionate number of internationally collected specimens (Fig. 2B), and ii) tend to have self-collected most of the specimens coming from their own countries (Fig. 2C). Further, over 80% of the specimens with digital images are held by European and North American institutions, the majority of which were collected from those two continents (Supplementary Fig. 3). We note that not all countries in these two continents have actively colonized others, but some have nonetheless amassed sizable international plant collections (e.g., Switzerland). Moreover, not all digitized specimen data online are available from GBIF – unique data can be found in smaller, regional repositories or institutional databases. Also, such online databases can harbor gaps and biases. Thus, the availability of digital data assembled for this study may be entirely reflective of the complete distribution of specimens collected and stored across the world ^31,32^. Nonetheless, these results are based upon the single largest curated digital specimen dataset currently available and represent our best estimates. To address these inherent limitations of our assessment of digitized specimen content we examined the distribution of specimens within physical herbaria across the world.

**Figure 2.**
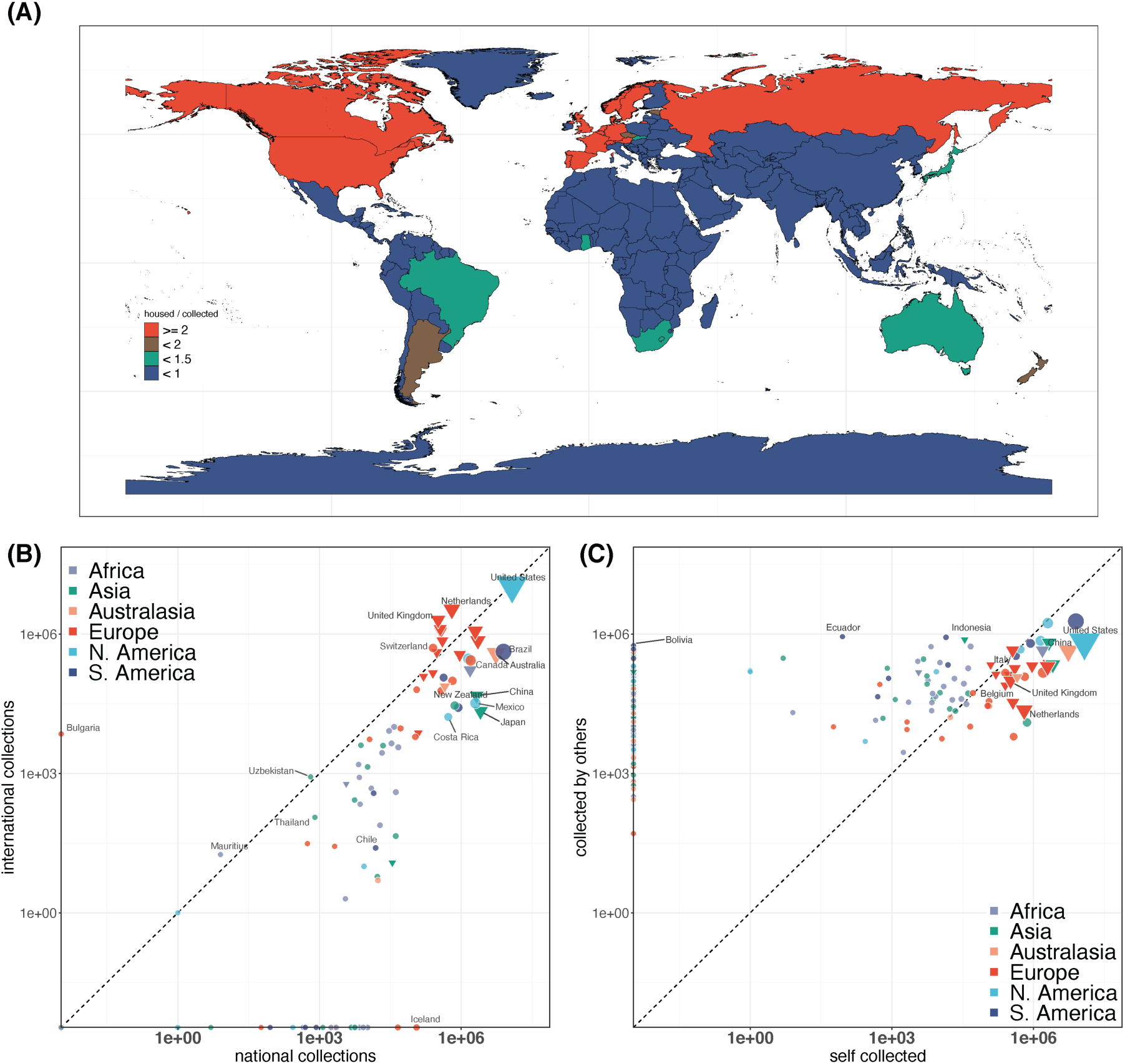
Disparity in the collection and housing of plant diversity. The top panel (A) maps the ratio of total number of species with specimens housed in a country to the total number of species collected in that country (species housed/collected). Ratios below one (blue) indicate areas where the number of species that have been collected from that country is higher than the number of species housed in that country. Panel (B) shows the ratio of national versus international collections held by each country, while (C) depicts the ratio of self-collected specimens in each country versus those collected by other countries. Points sizes are log-scaled to the total number of specimens. Triangles represent countries that have overtly colonized other countries in the past following reference ^29^.

### A glimpse inside the cabinet

Increasing digitization of specimen data and their online mobilization have seemingly decentralized and democratized access to herbarium data greatly ^33^. As demonstrated above, open access biodiversity data repositories such as GBIF and iDigBio allow researchers from around the world to query aggregated specimen metadata and images, alleviating some of the need for extensive and prohibitive travels to consult materials and requests for loans. Institutional databases, although containing fewer specimens than global databases, efficiently contribute to make available their own holdings, and encourage worldwide researchers to request free high-resolution images and better define loan requests. However, digitization requires significant investments in infrastructure (i.e., physical space, photographic devices, data storage) and personnel, which is often not financially feasible for smaller institutions and developing countries. Along these lines, it has been argued that digitization could exacerbate the exploitation of intellectual property and biological resources by developed nations in a form of neo-imperialism ^34^. Further, only a small portion of specimen data are digitized and shared online at this point in time, and there are many studies that require access to physical specimens.

According to *Index Herbariorum* ^1^, there are at least 3426 herbaria globally that together house approximately 400 million specimens. Over 60% of these herbaria and 70% of specimens are located in developed countries with colonial histories (Supplementary Fig. 4). To better understand the current state of the world’s collections and their digitization, we conducted a survey of major herbaria as listed by *Index Herbariorum* and targeted representative regional herbaria. A total of 93 herbaria across 39 countries and 6 continents submitted at least partial responses to the survey that could be used. Similar to the patterns observed using digitized data from GBIF, we found that herbaria in developed nations with colonial histories in North America and Europe housed a higher proportion of internationally collected specimens on average (Fig. 3A). This pattern generally held consistent across databased specimens with collection date and location information (Fig. 3B) and specimens with digital images (Fig. 3C) shared online. There were some exceptions; for instance, herbaria in Singapore hold a disproportionate volume of international collections, possibly due to its small size, location, history as the main British colonial outpost in the area, and past and present association with Malaysia.

**Figure 3.**
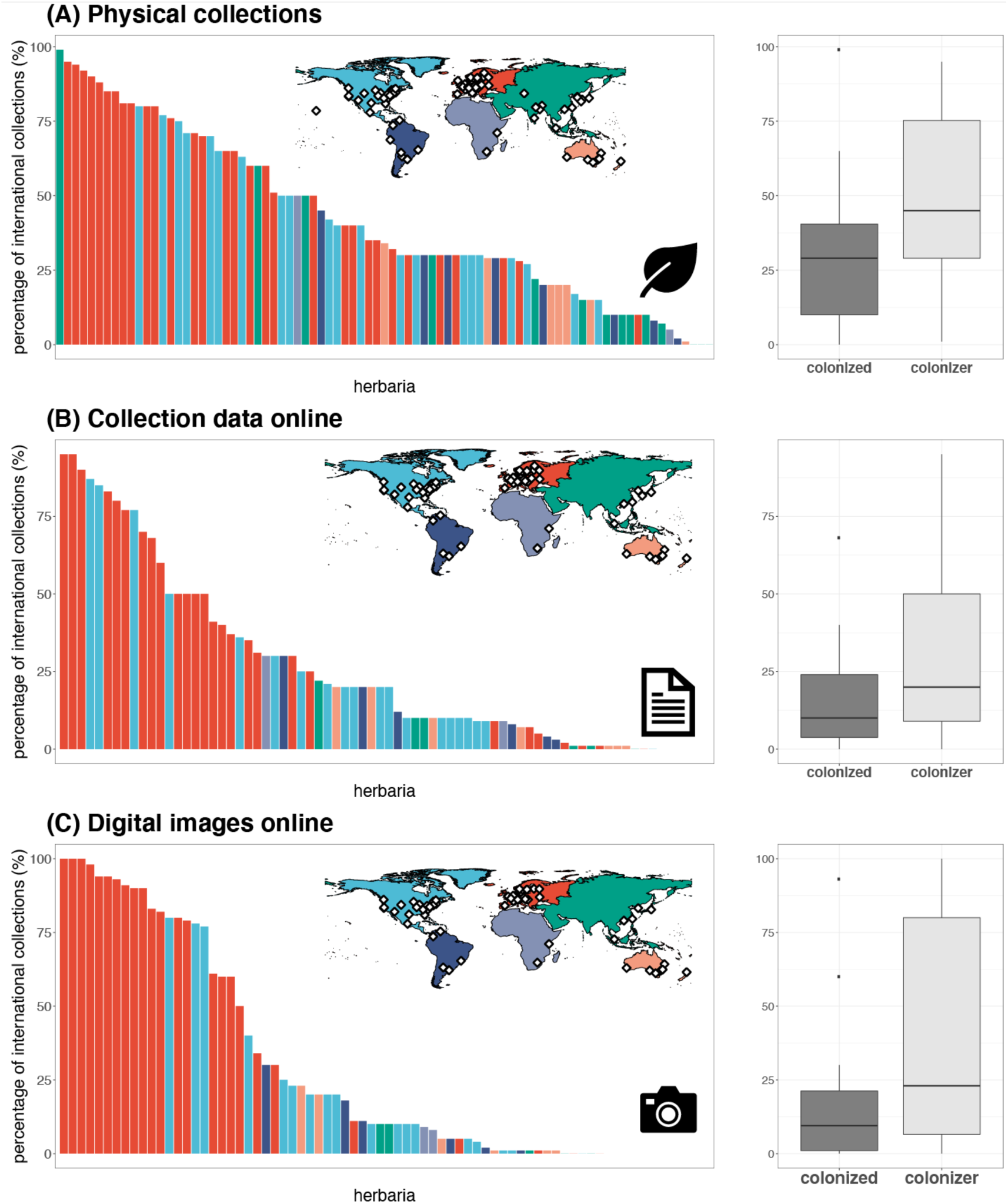
The percentage of internationally collected specimens in herbaria. Trends across physical specimens (A), specimens with at least a portion of their metadata available online (B), and specimens with digital images shared online (C) are illustrated as histograms where each bar represents a surveyed institution, and colors indicate different continents. Boxplots to the right summarize this information among countries that have been colonized, versus those that have colonized others following reference^29^. Countries that both experienced colonization and colonized others are depicted under their most recent category.

Our surveys also revealed that the digitization of herbarium specimens remains in its infancy. We estimated that overall, less than 30% of physical collections have at least collection location and date information online, and less than 10% have available digital images (Fig. 4A). Nearly all surveyed herbaria have ongoing digitization efforts with at least some specimen data provided online (Fig. 4B-C). However, these data are not always widely accessible, and represent only the tip of the iceberg relative to the physical collections and are thus woefully insufficient to address the reverse-latitudinal gradient of diversity inside herbarium cabinets. Our results suggest that the patterns we observe from GBIF data are likely representative of the larger reserves of specimen data yet to be digitized and mobilized online. Indeed, most institutions gave equal priority to the digitization of national and international collections (Fig. 4D) and submit their digitized specimen data to GBIF and/or regional databases that often share data with GBIF (e.g., Consortium of California Herbaria, the Australasian Virtual Herbarium, eReColNat, Virtual Herbaria - JACQ; Fig. 4E). Other aggregators such as JSTOR - Global Plants and iDigBio share data with GBIF as well ^35^. Our survey results also highlight the fact that we are still in the infancy of digitizing herbaria, and thus retain the opportunity to reassess how ongoing and future digitization and mobilization efforts can be organized to better address the colonial legacy of these collections.

**Figure 4.**
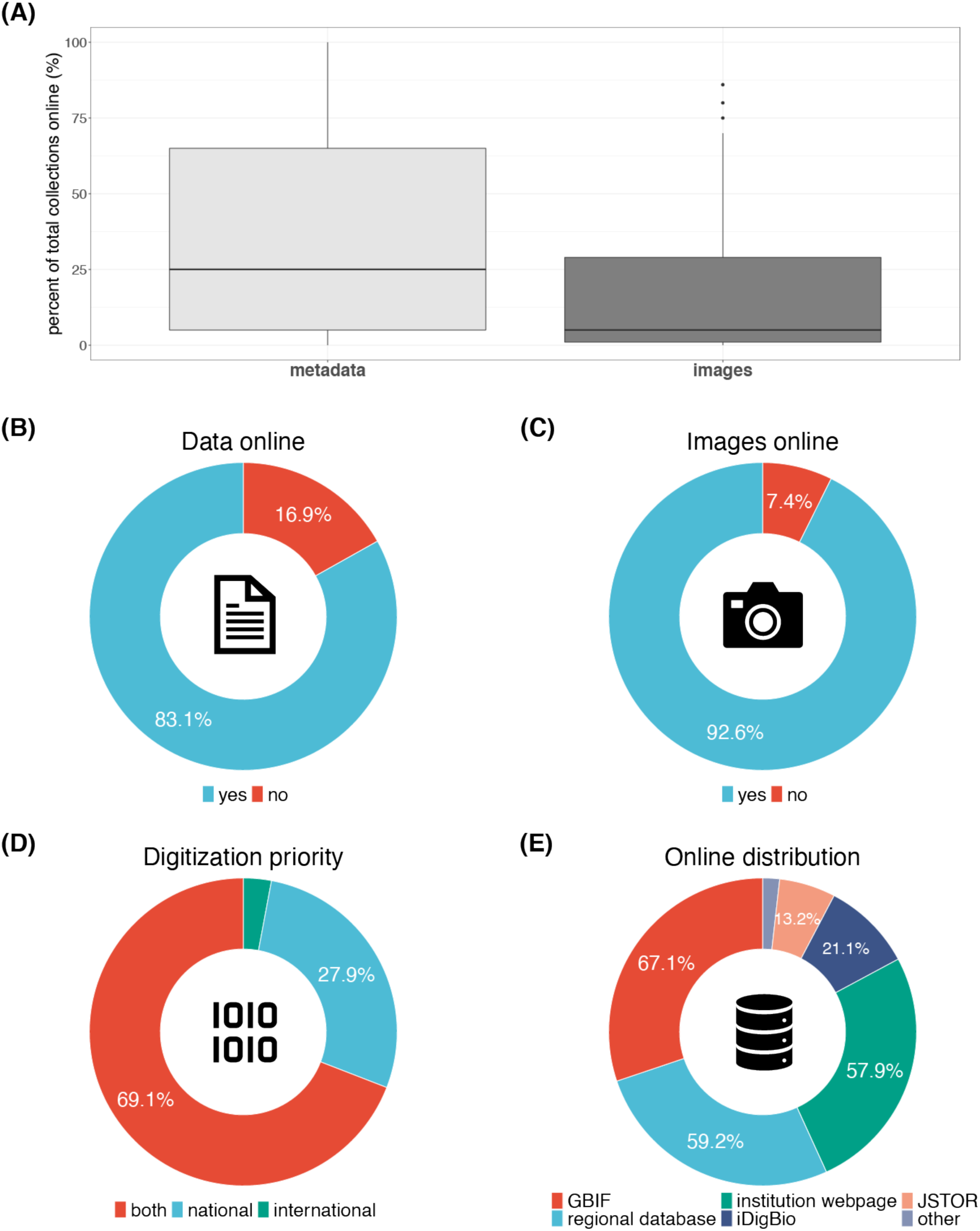
Trends in the digitization of herbarium specimens. The boxplot (A) summarizes the percentage of physical collections in surveyed herbaria that have at least location and date information (metadata) or digital images available online. Pie charts (B) and (C) show the percentage of herbaria that have at least some data and images of their collections shared online, respectively. The digitization priority of herbaria in terms of where specimens were collected is illustrated in pie chart (D), while pie chart (E) shows how the surveyed herbaria share and distribute digital specimen data. Multiple answers were possible in panel (E).

### Challenges and opportunities

Our study demonstrates a major disparity between where plant diversity naturally exists and where it is artificially housed and catalogued. This renders much of the world reliant on botanical knowledge and resources housed outside of their own borders. This disparity not only impacts capacity for conservation and basic research, but commercial and government enterprises that seek to appropriate and monetize biological resources and their derivatives as well. In addressing this disparity, recent discussions regarding approaches to decolonize cultural institutions, natural history museums, and biogeographical practices in general must be applied to herbaria as well.

First, as Das & Lowe (2018) note ^36^, it is important to acknowledge the colonial legacy of herbarium collections and to present the history and circumstance of these collections alongside existing interpretations about the specimens and their role in scientific research. They argue that such acknowledgment is a critical step towards bridging the gap between natural history collections and audiences in previously colonized nations and ensuring inclusiveness in the collection, curation, and use of these collections. One way to openly share and communicate such narratives is via themed exhibitions and tours, such as the black history tours of Hintze Hall at London’s Natural History Museum or the First Nations-led and informed “Unsettled” exhibition at the Australian Museum. These tours recognize and emphasize the (unrecognized) contributions of Indigenous peoples to the culture, science, and natural history on display. Though most herbaria traditionally do not offer public exhibitions and herbarium specimens are rarely prominent in natural history museum displays (in part, due to their fragility), increasing specimen digitization efforts have made it possible to curate digital exhibitions and virtual tours without competing for space and attention with others considered more charismatic (e.g., large mammals and dinosaurs). Awareness and acknowledgement can also be facilitated by including positionality statements in grant proposals, research articles and other scientific communications that involve herbarium collections. Positionality statements describe the position of a researcher in relation to the social and political context of all phases of the research in question and are well-established in the humanities but still rare in the natural sciences ^37^.

Second, we must continue to improve accessibility to the vast information held in herbaria worldwide, for both scientists and the public. Though digitizing and sharing specimen data is hardly a new idea, our survey suggests that the data currently available only represent a small portion of what resides inside herbarium cabinets. Indeed, several of our survey participants noted that estimating the size and distribution of their collections is difficult – only a small portion of herbarium specimens have been databased by their respective institutions, and no formal inventories exist. Though massive digitization efforts have been funded, particularly at institutions in developed countries, it is ironic that even these institutions often lack the funding for adequate curation and processing of specimens. Our analysis of available digital collection data also shows that higher-level data products (i.e., images) for many previously colonized areas are lacking (Supplementary Fig. 3). Digitization efforts focused on increasing representation from such areas could help bridge the reverse-latitudinal gradient of plant diversity knowledge. Further, though much of the data that has been digitized from herbarium collections are shared via open data repositories (e.g., GBIF, iDigBio, BIEN, SpeciesLink, AVH, eReColNat), a large portion remain available only upon request (which can be denied), paywalled, or inaccessible outside of specific groups. Targeted initiatives and funding opportunities that prioritize the curation, digitization, and sharing of collections from developing countries can be one way to address these discrepancies. There have been some promising efforts along these lines, such as the NSF supported GLOBAL Bryophyte & Lichen Thematic Collections Network, and the Mellon Foundation’s African Plants Initiative ^38^. We can also increase support for loan and exchange programs across herbaria, facilitating access and repatriation of physical specimens as well. Such efforts must be mindful of the legacy of some herbarium collections. For instance, specimen returns in accordance with permits or agreements are traditionally referred to as “gifts”, but it may be preferable to use a different term, such as “returns” ^39^. Also, we must be mindful that specimens can contain biocultural information that is culturally inappropriate for broader circulation, and can risk further exploitation of Indigenous cultural knowledge. Thus, efforts to improve accessibility to botanical collections and share knowledge therein require careful discourse with all parties.

Third, in in addition to recognizing the sovereignty of a nation’s biological resources and that biodiversity can be best studied where it occurs ^40^, capacity-building in previously colonized countries through the sharing of tools and knowledge for contributing towards research is critical – if the science resulting from collections is globally relevant, the means of contributing should be distributed as such ^37,41^. In particular, it is crucial to ensure that local contributions are sufficiently recognized and facilitate the development of local research priorities and agendas during this process. Acknowledging the providing country personnel in all aspects, from specimen labels to publication authorship to grant proposals is critical. Further, the digital products of herbarium specimen data could be hosted and managed by researchers in the countries where they were originally collected as a form of repatriation, who could be trained and supported as necessary by institutions with greater capacity. Although the latter might not necessarily dispose of the necessary funding to support the local partner, they could play a major role when a grant request is addressed to an international agency, clearly stating their engagement in the transfer of technical and scientific knowledge. International collectors should be mindful to leave duplicate specimens in the host country – this practice has become increasingly adopted over recent decades, and at times enforced by local governments, especially since the Convention on Biological Diversity was signed in 1992 and the Nagoya Protocol on Access and Benefit Sharing was drafted in 2010 (https://www.cbd.int/abs/). Still, many regions lack the facilities to store and curate collected specimens. In such cases, collectors could gather and treat duplicate specimens as loans until the necessary local infrastructure is established. This would in turn facilitate a more equitable, global view towards the collection, curation, and use of herbarium specimens. To support such efforts, we strongly recommend that grant proposals involving the collection/curation/digitization of specimens associated with developing countries include requests for funding to support local colleagues and collaborators. Institutions and scientists need to seek ways to expand opportunities for partners in providing countries to participate in research design and grant application, in addition to activities directly pertaining to the collection and curation of specimens. In turn, funding bodies must recognize the need to support local partners appropriately and guarantee access to the knowledge and benefits arising from plant collections sampled abroad. Importantly, these and other efforts to decolonize herbaria should be guided by the needs and wishes of previously colonized peoples. One example of such a partnership can be found in a recent project to sequence and study the genome of the tuatara, a cultural treasure of the Māori people ^42^. The Indigenous peoples provided access to the species and associated knowledge, and were involved in all decision-making regarding the use of the genomic data generated by the study and any benefits that may accrue.

A profound set of challenges lie ahead if we are to address the still-persistent legacy of colonialism in our plant collections. However, ongoing digitization efforts have offered us new avenues of deploying knowledge and infrastructure and sharing the benefits arising from the utilization of herbarium collections. Science is not exempt from sociopolitical realities and we should not avert our gaze from the inconvenient origins of these otherwise precious resources. To this end, we have endeavored to provide a glimpse into the extent of the colonial legacy that plagues our herbarium collections. Only by embracing these realities can we progress towards a more inclusive global herbarium.

## Acknowledgements

We would like to acknowledge the past and continuing contributions of colonized peoples to botanical science and knowledge. We also thank the herbaria that contributed data to this work: AAU, AD, ASSAM, ASU, B, BISH, BM, BO, BP, BR, BRI, BRNU, BSA, BSHC, BSID, C, CANB, CAS, CHR, CL, CM, COL, CORD, DAO, DUKE, EA, F, FI, FLAS, FR, G, GB, GH, A, ECON, AMES, FH, NEBC, GJO, GOET, GZU, H, HAL, HIRO, IBSC, SI, K, KH, KW, L, LBL, LE, LIL, LY, M, MA, MEL, MEXU, MICH, MIN, MO, MT, MW, NCU, NIBR, NICH, NSW, NY, O, OULU, P, PC, PAD, PE, PERTH, PRC, PRE, NBG, NH, QFA, RM, USFS, RO, S, SI, SING, SP, STR, TAIF, TASH, TENN, TEX, LL, TNS, TUR, UC, JEPS, UPS, US, USM, VEN, W, WTU, ZT. DM was supported by AAAA-A19-119031290052-1 “Vascular plants of Eurasia: systematics, flora, plant resources”.

## Author contributions

D.S.P. conceived the initial idea with C.C.D. which was refined through discussions with X.F. D.S.P and X.F. analyzed specimen data and D.S.P. designed the survey with C.C.D. D.S.P. supervised the study and wrote the original draft with input from X.F. All other authors provided data from their respective herbaria. All authors contributed to further revising the manuscript.

## Data availability

Data discussed in the paper are either publicly available through GBIF (https://www.gbif.org/), *Index Herbariorum* (http://sweetgum.nybg.org/science/ih/), or attached supplements.

## Supplementary Information

### Methods

We downloaded plant specimen data (kingdom = Plantae; basis of record = preserved specimen) from GBIF on April 23, 2021 (https://doi.org/10.15468/dl.nt5wkx). We only kept specimen records with accepted scientific names, valid country code and publishing country names. With the remaining 50,303,354 records, we compiled a country-by-country matrix that summarized the number of specimens collected from one country and housed in another. The country where a specimen was collected was based on the field “countryCode” and the country where a specimen was housed was based on the field “publishingCountry”. We also grouped the country-by-country matrix into a continent-by-continent matrix. To examine the temporal trends of collection, we further examined the data after separating them into two subsets; before and after 1945, which marks the end of World War II and the era of overt colonialism. We finally verified our analyses on a subset of data that i) had coordinates; ii) had the “countryCode” field matching the location inferred from the coordinates; and iii) were determined to be without geospatial issues by GBIF.

As records on GBIF represent a subset of the collections in herbaria across the world, we expanded our investigations to physical institutions. We sent out a survey in 2020 to major herbaria across the world as listed by *Index Herbariorum* (http://sweetgum.nybg.org/science/ih/) and select representative regional herbaria. Questions were focused on identifying the size of the collections, where they were collected, and the proportion digitized. A total of 93 herbaria across 39 countries and 6 continents submitted at least partial responses to the survey (Supplementary Data 1).

We recognize that certain assumptions were made in these analyses. First, the Western scientific system is not the only way to understand and describe botanical knowledge, and though many of our discussions pertain to such as it is broadly adopted, we do not mean to devalue or reject other knowledge systems. Second, we use geopolitical constructs that are not free from the influence of colonialism. For instance, though we treat Australia as a single entity, it is home to over 500 Aboriginal nations. Finally, though we posit that the era of overt colonialism has ended, we realize that there was no single process of decolonization, and that the idea that colonization is over can be problematic as its legacy persists to this day, even in botanical collections. Along these lines, here we use the term colonization in a fairly general sense to describe a relationship between two countries, independently of their level of development, in which one has subjugated and governed the other over a period of time, contributing to the current state of its institutions following reference ^29^.

**Supplementary Figure 1.**
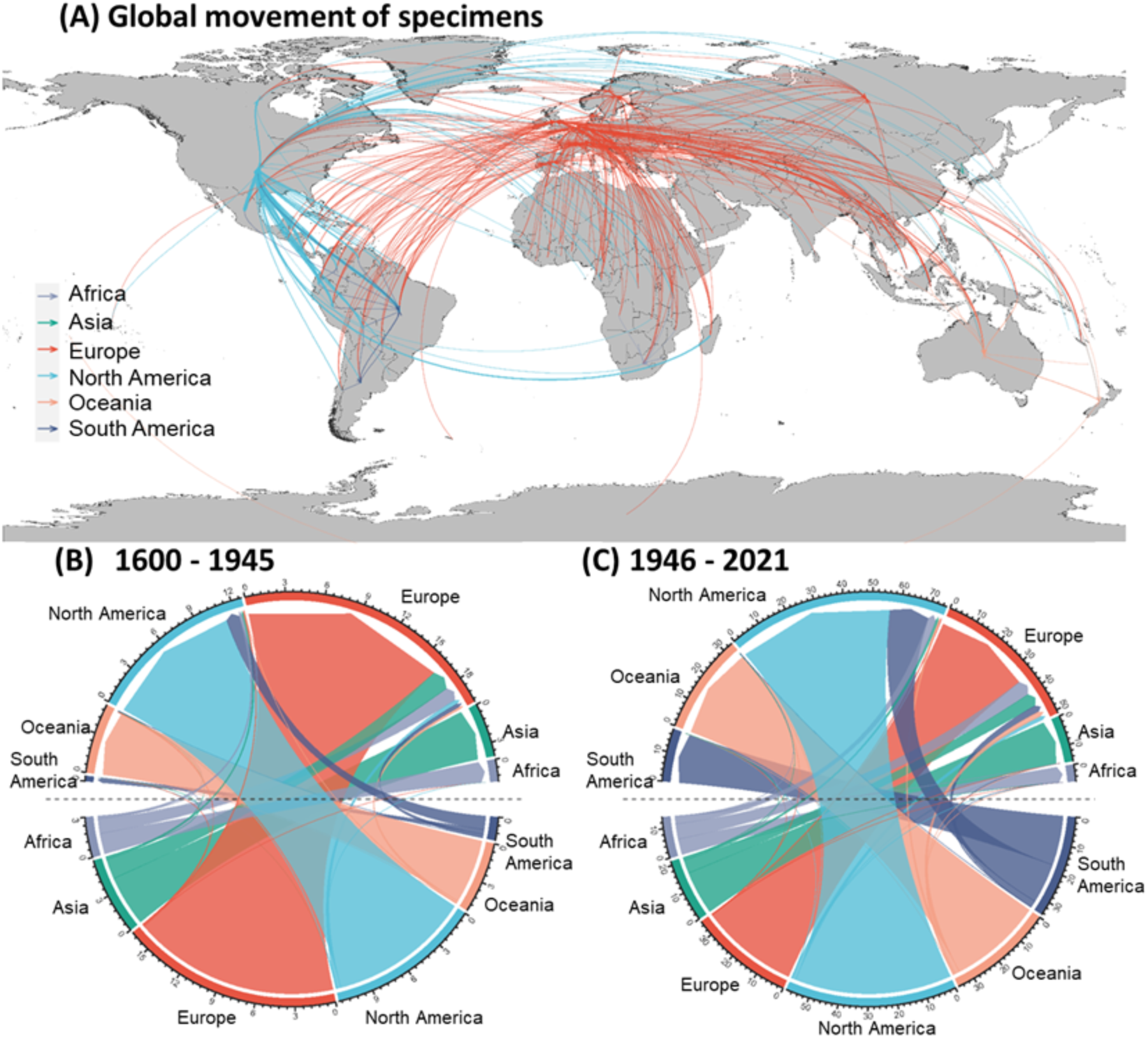
The past movement of plant specimens across the globe based on GBIF records with geographic coordinates. The world map depicts the top 10 percentile of intercontinental connections between countries where specimens have been collected and where they are currently housed regardless of collection date (A). The widths of the arrows are proportionate to the number of specimens dislocated and are colored by destination continent. Collections that remained in the country of collection are not depicted. The lower panels illustrate the intercontinental flow of specimens before (B) and after (C) the end of overt colonialism post World War II (late-1945). Arrows are colored by the continent of origin. Numbers on the outer ring indicate specimen numbers collected from (lower half) or stored in (upper half) each continent and are in multiples of 100,000. Colors on the outer ring represent different continents.

**Supplementary Figure 2.**
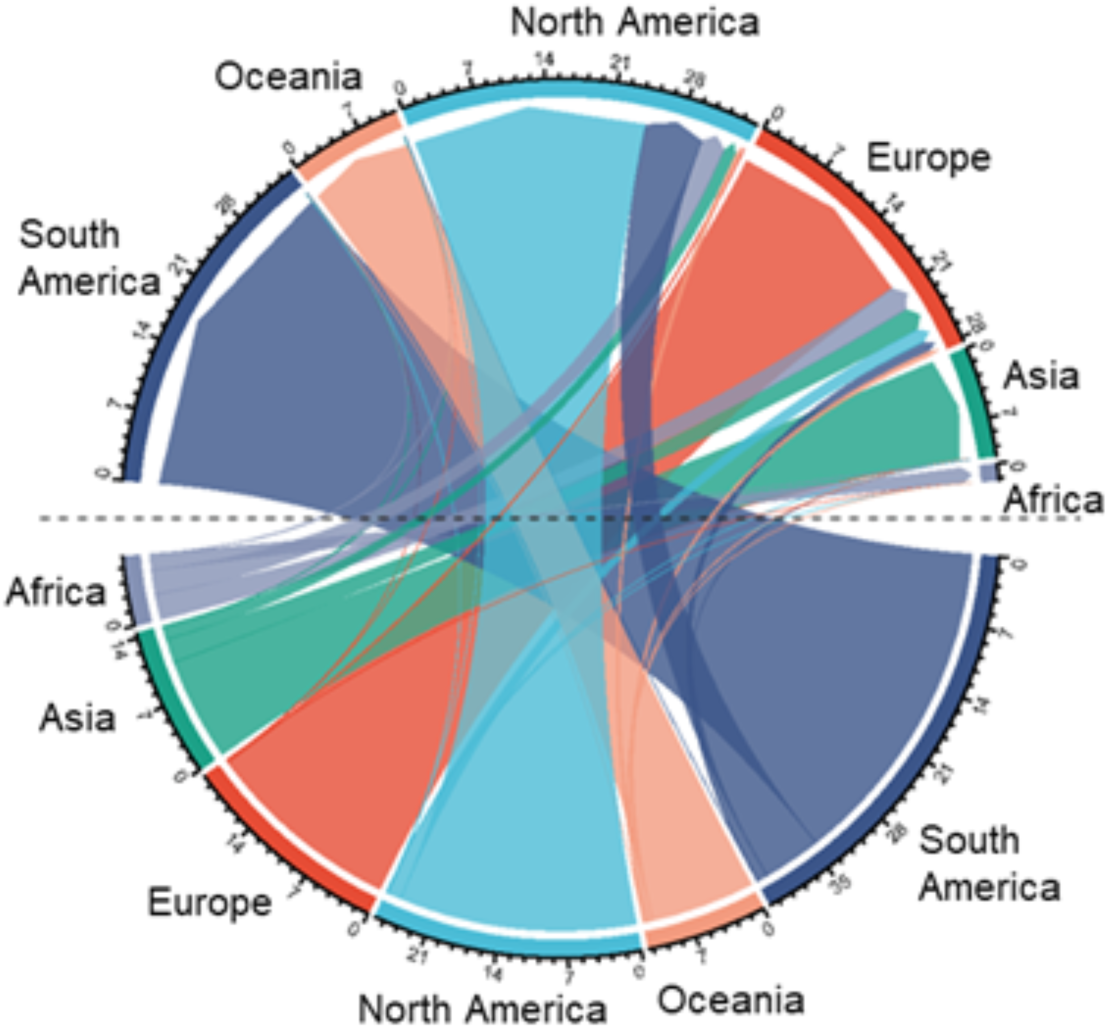
Global origins of specimens collected in the 21st century. The network illustrates the intercontinental flow of specimens. Arrows are colored by the continent of origin. Numbers on the outer ring indicate specimen numbers collected from (lower half) or stored in (upper half) each continent and are in multiples of 100,000. Colors on the outer ring represent different continents.

**Supplementary Figure 3.**
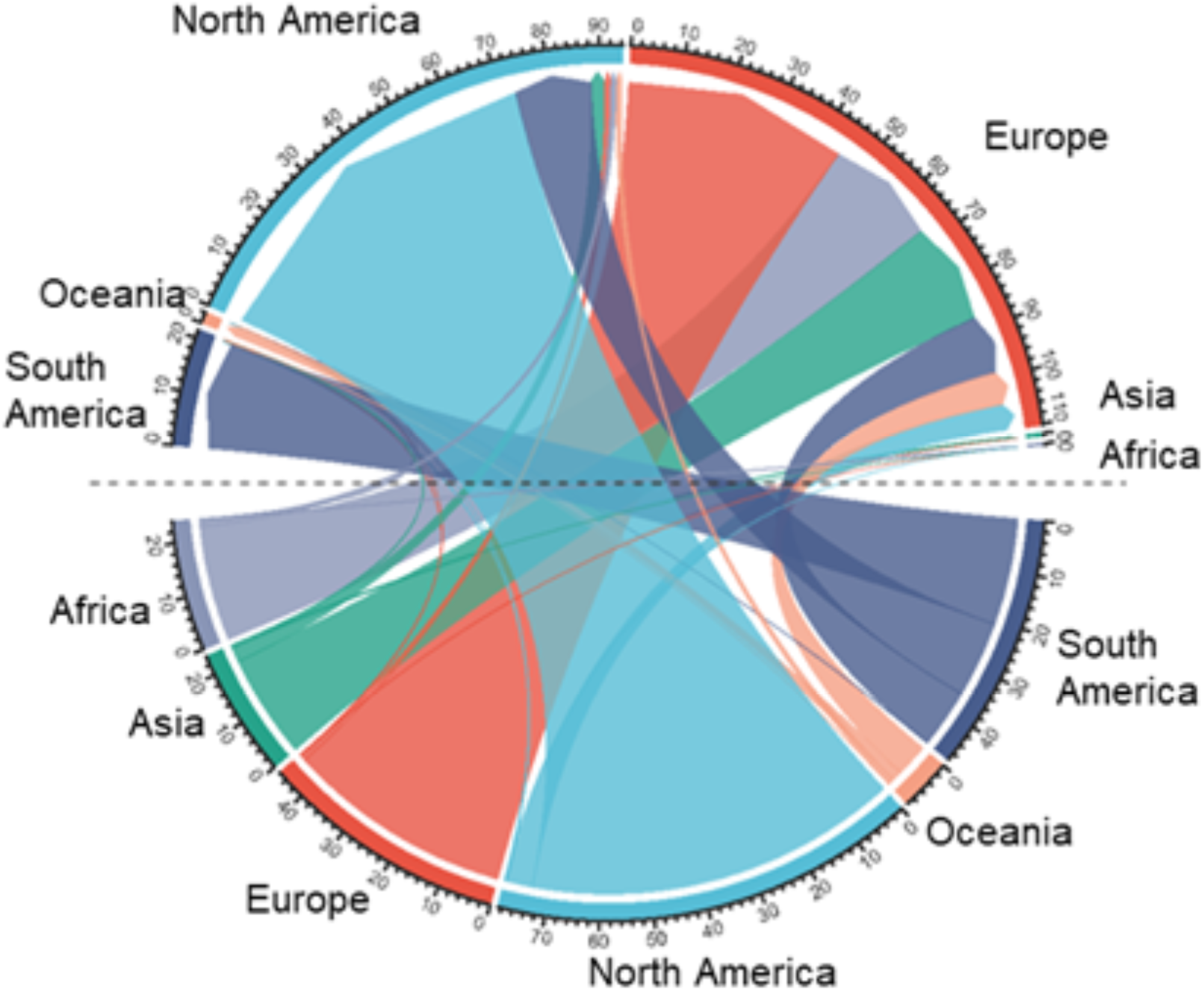
Global origins of specimens with digital image data. The network illustrates the intercontinental flow of specimens with digital images available. Arrows are colored by the continent of origin. Numbers on the outer ring indicate specimen numbers collected from (lower half) or stored in (upper half) each continent and are in multiples of 100,000. Colors on the outer ring represent different continents.

**Supplementary Figure 4.**
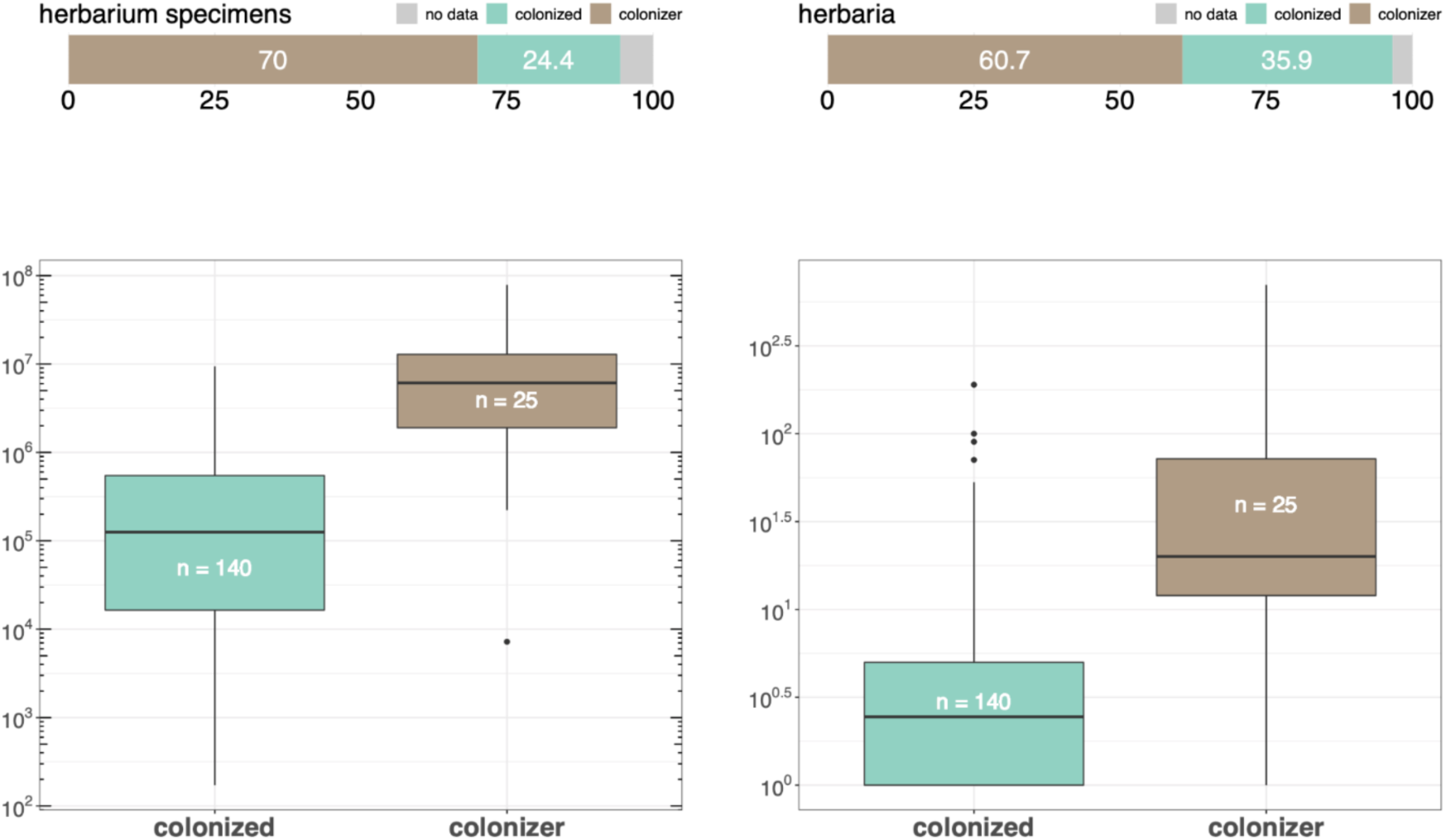
The distribution of herbaria and herbarium specimens across colonized and colonizing nations. Top panels depict the percentage of global herbaria and herbarium specimens situated in these nations. Lower panels contrast the number of herbaria and herbarium specimens held in institutions in colonized and colonizing nations. Countries that both experienced colonization and colonized others are depicted under their most recent category.

**Supplementary Data 1**. Survey results

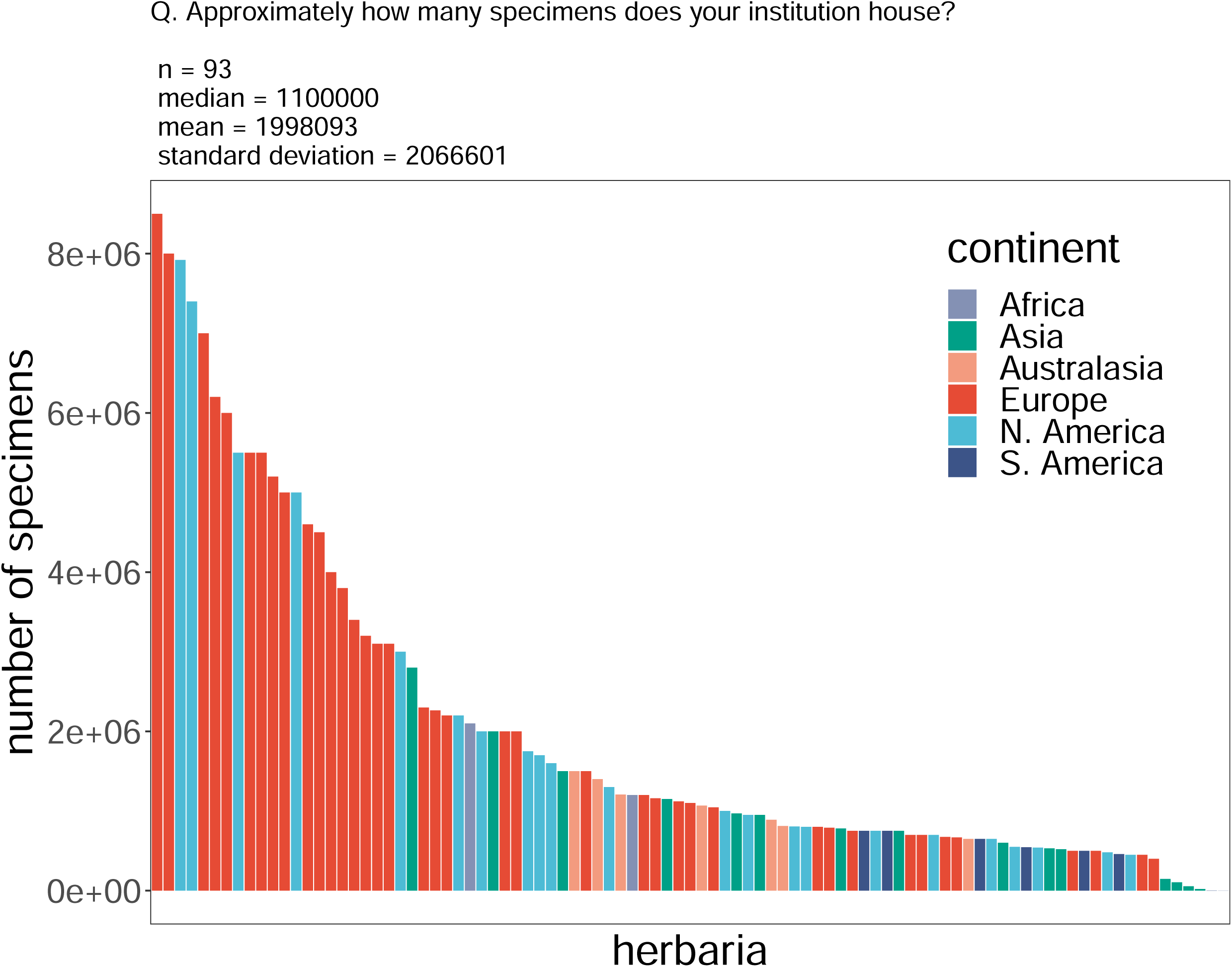

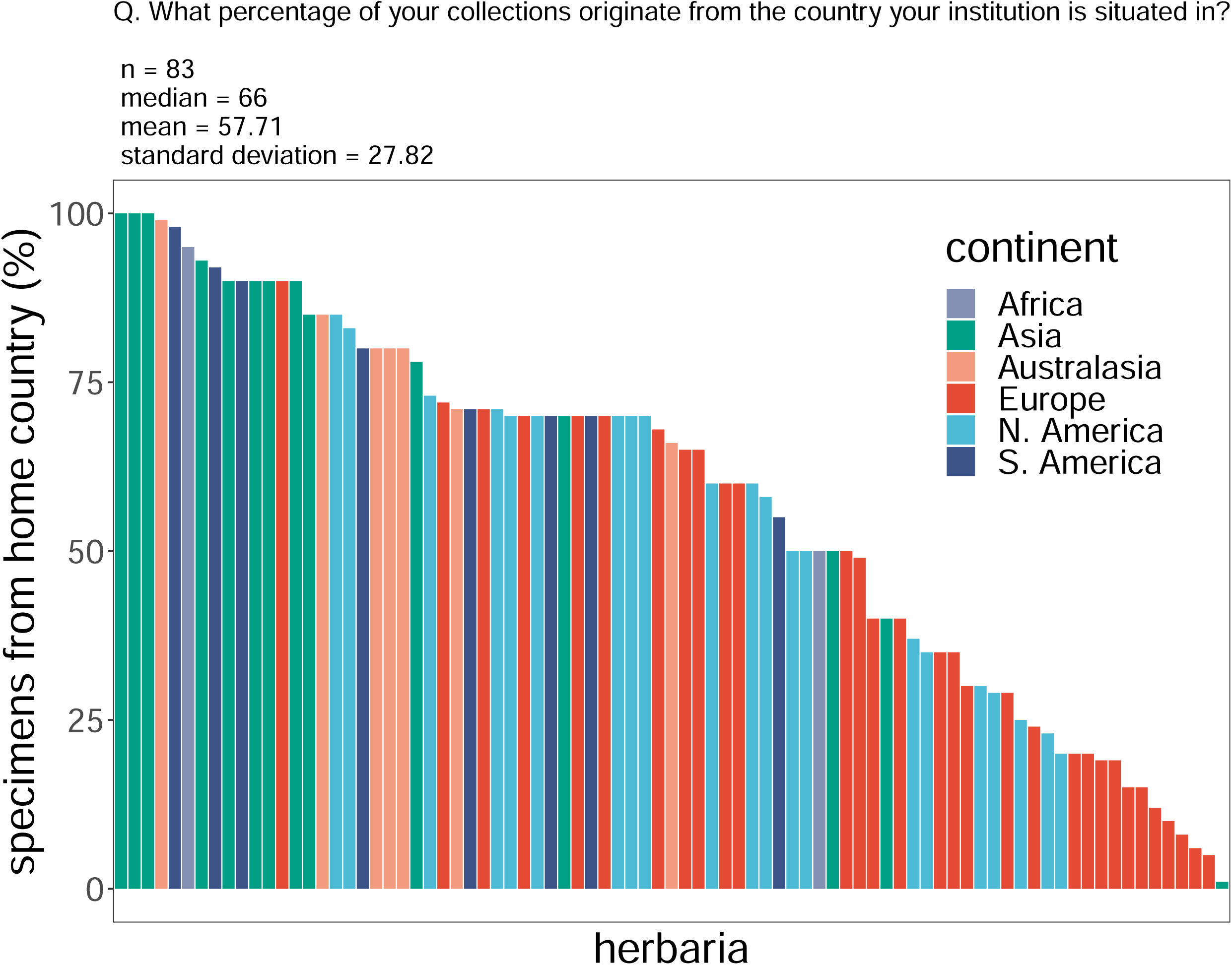

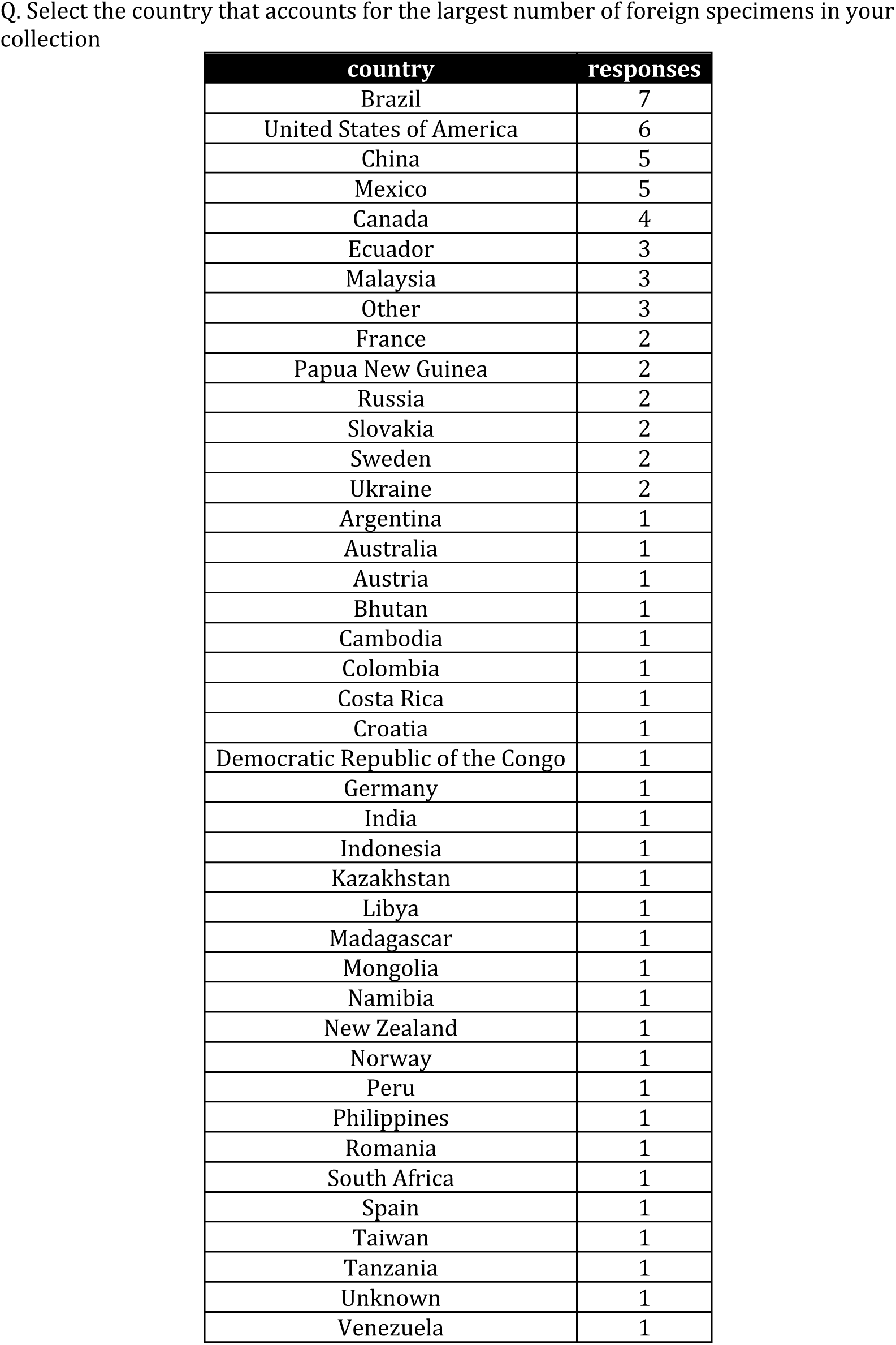

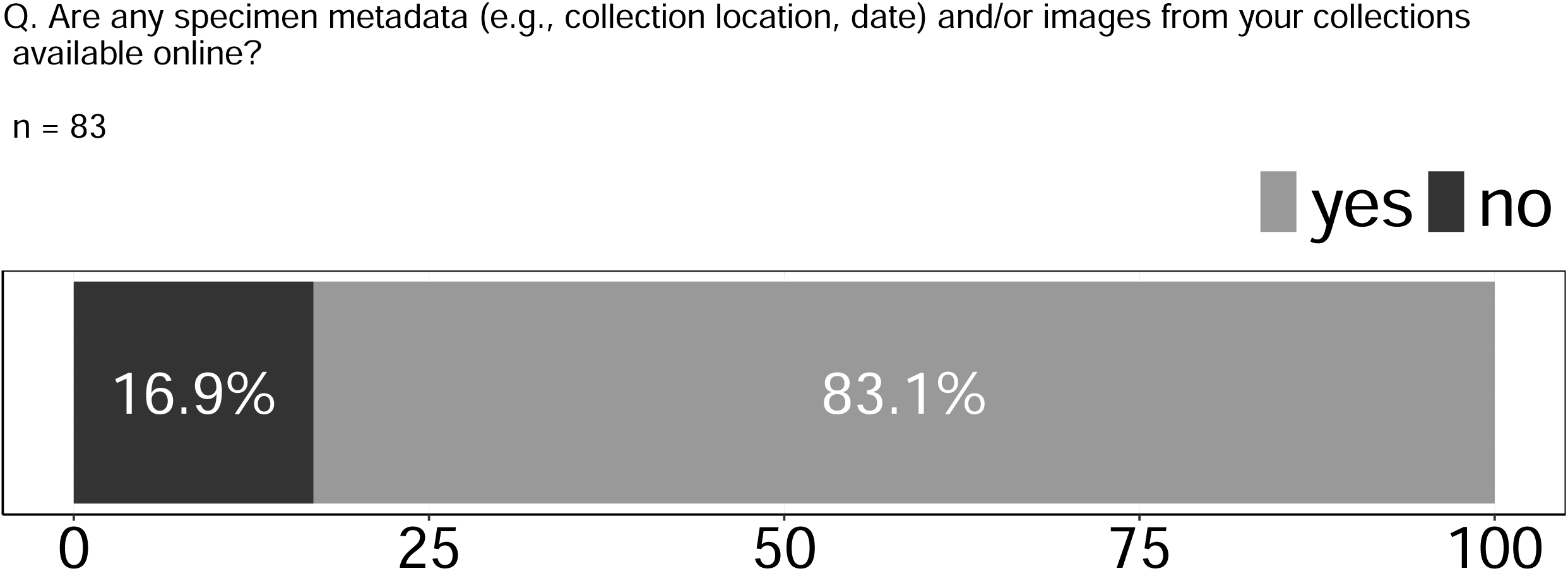

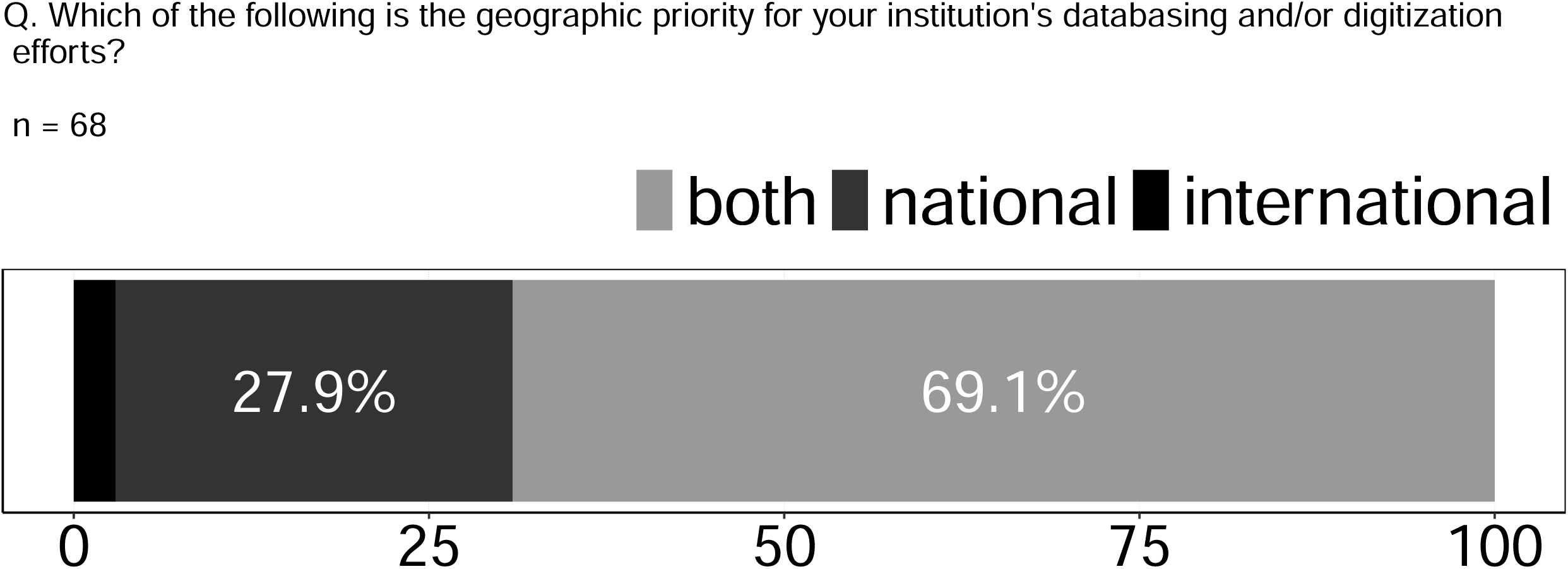

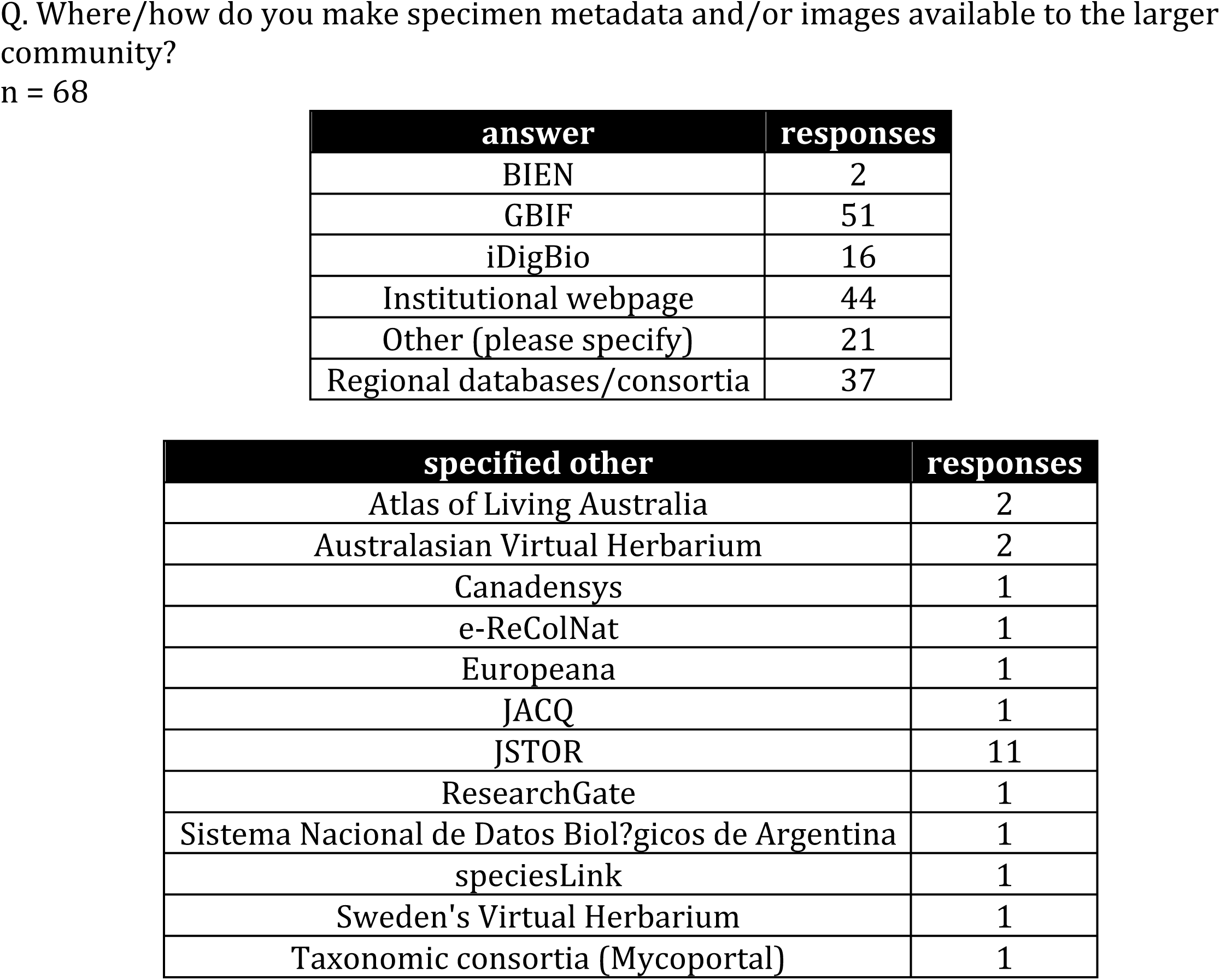

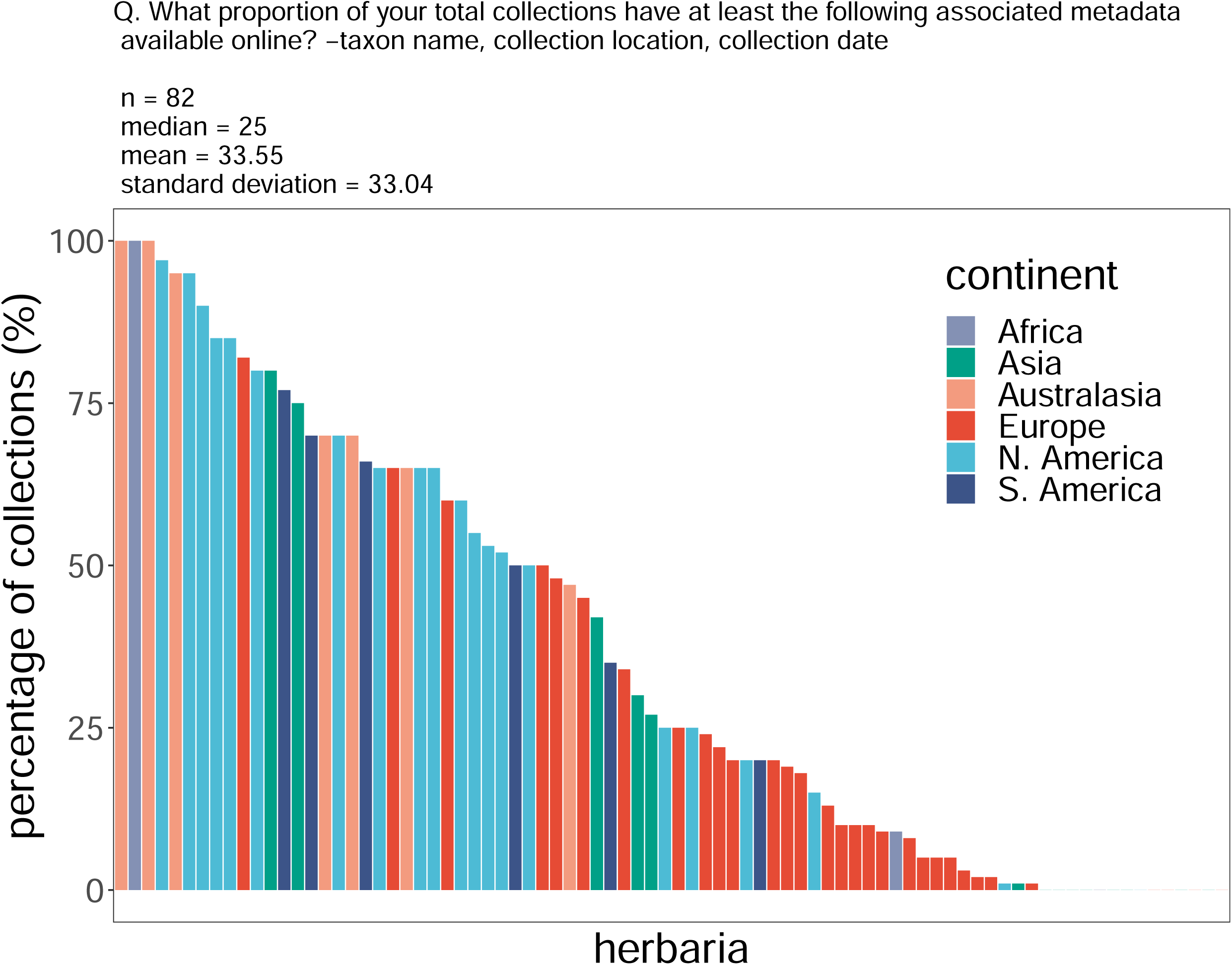

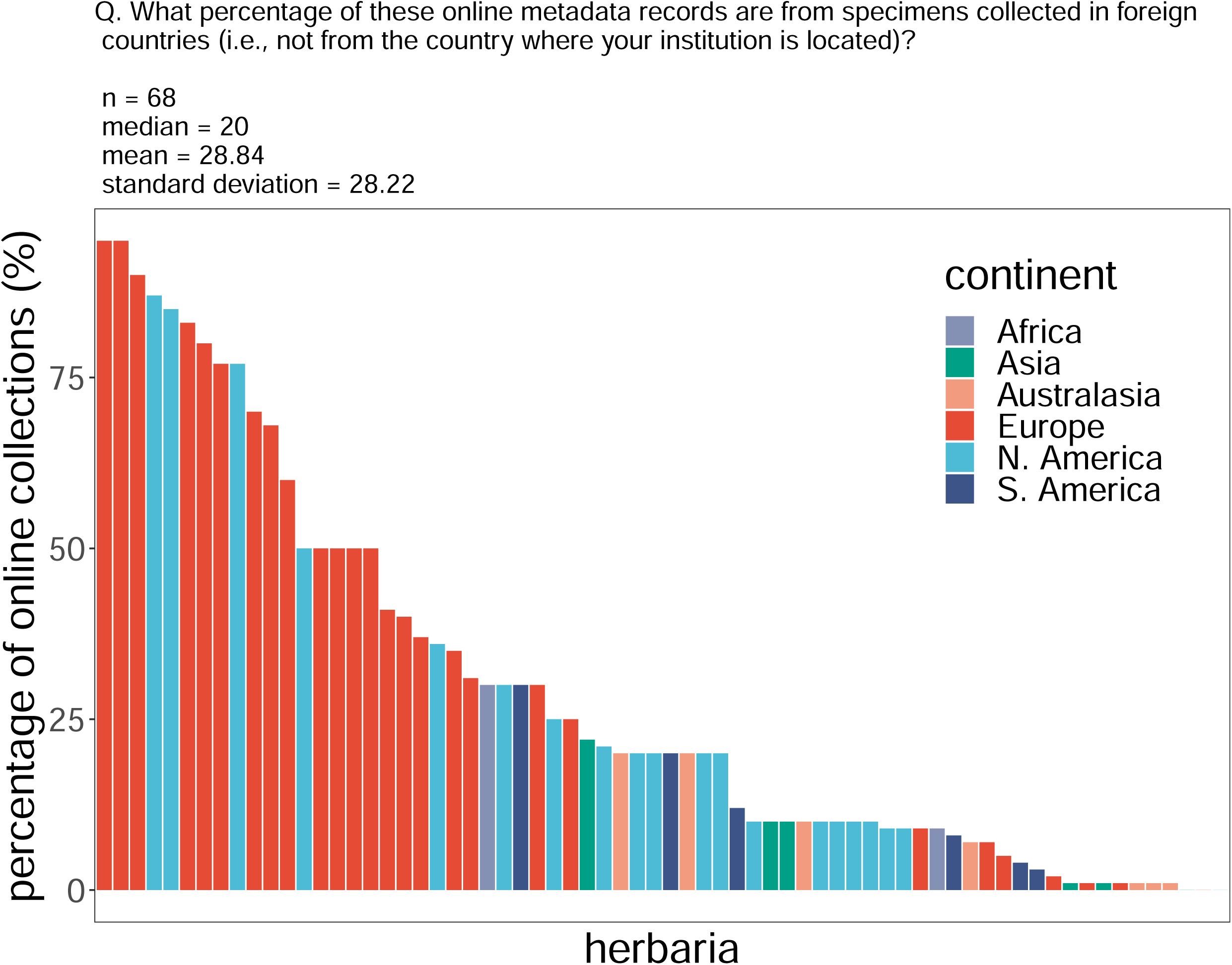

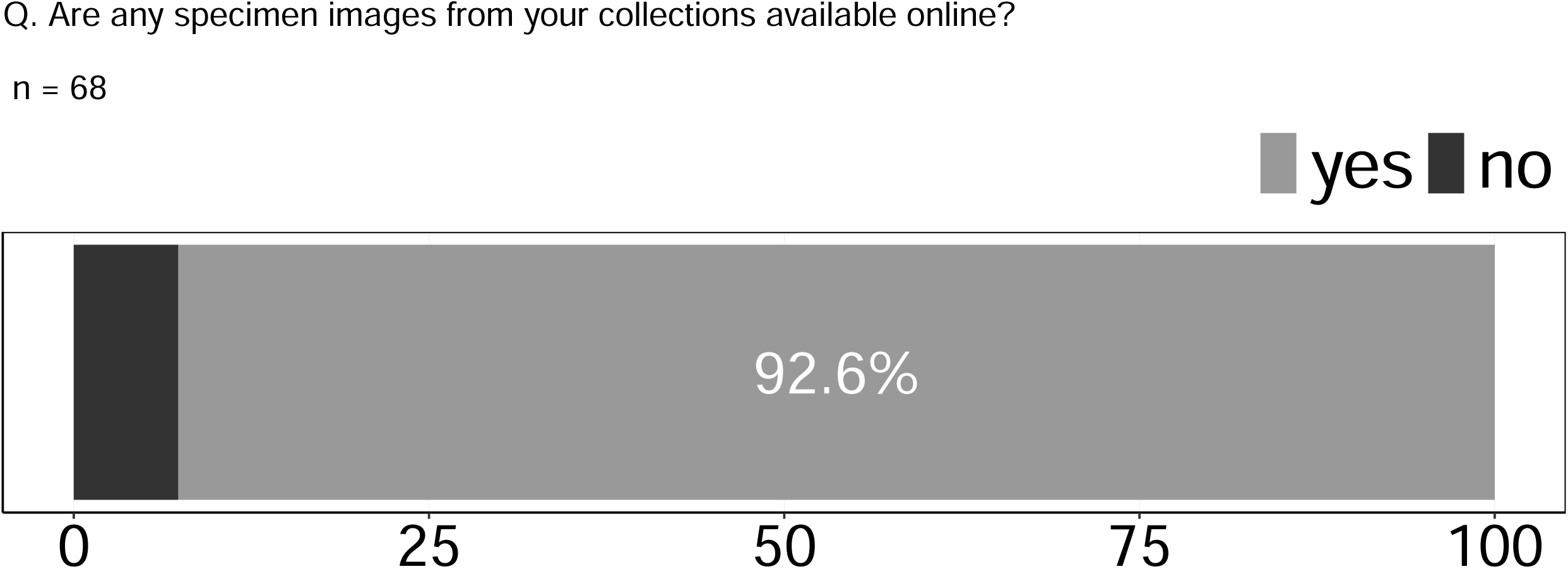

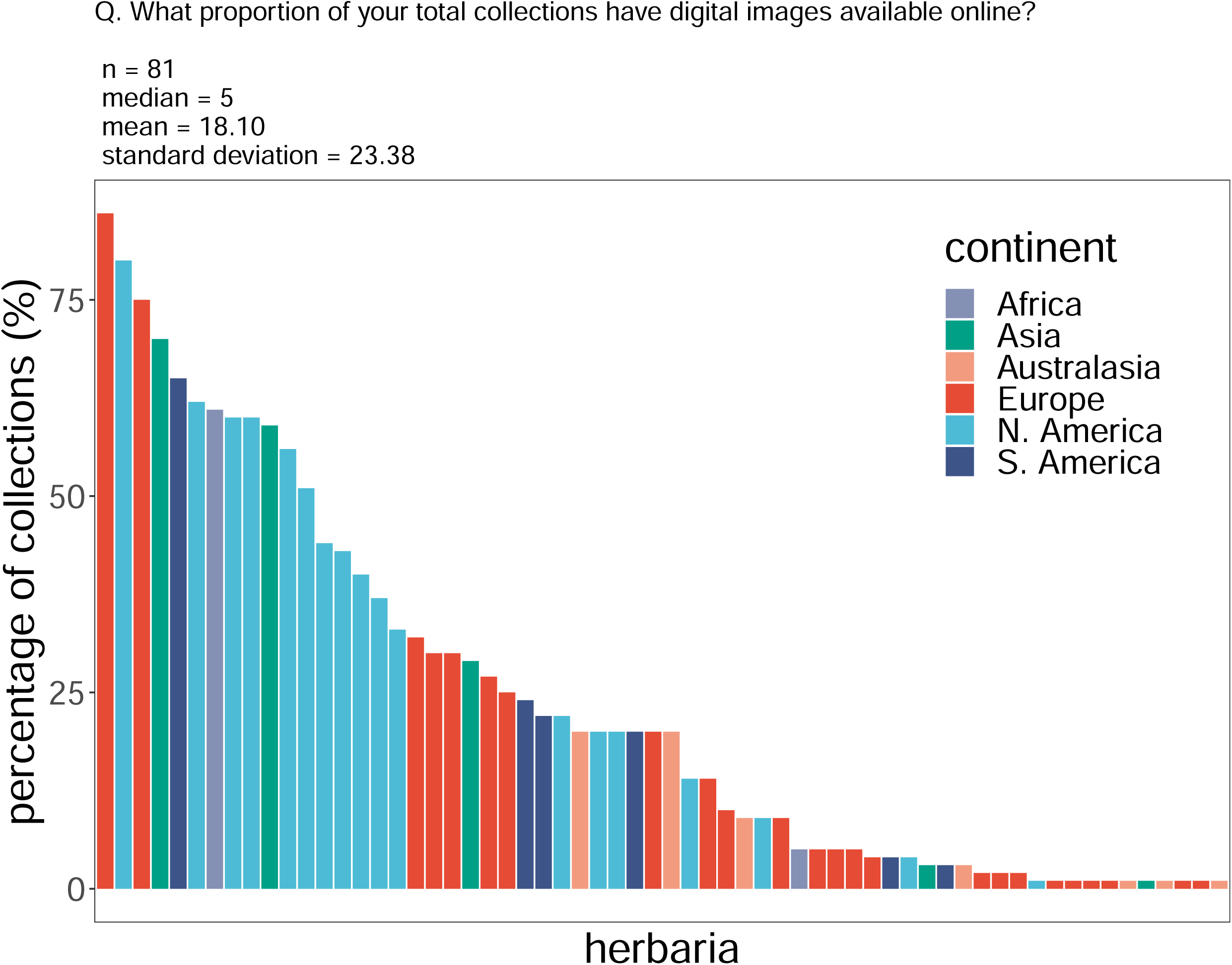

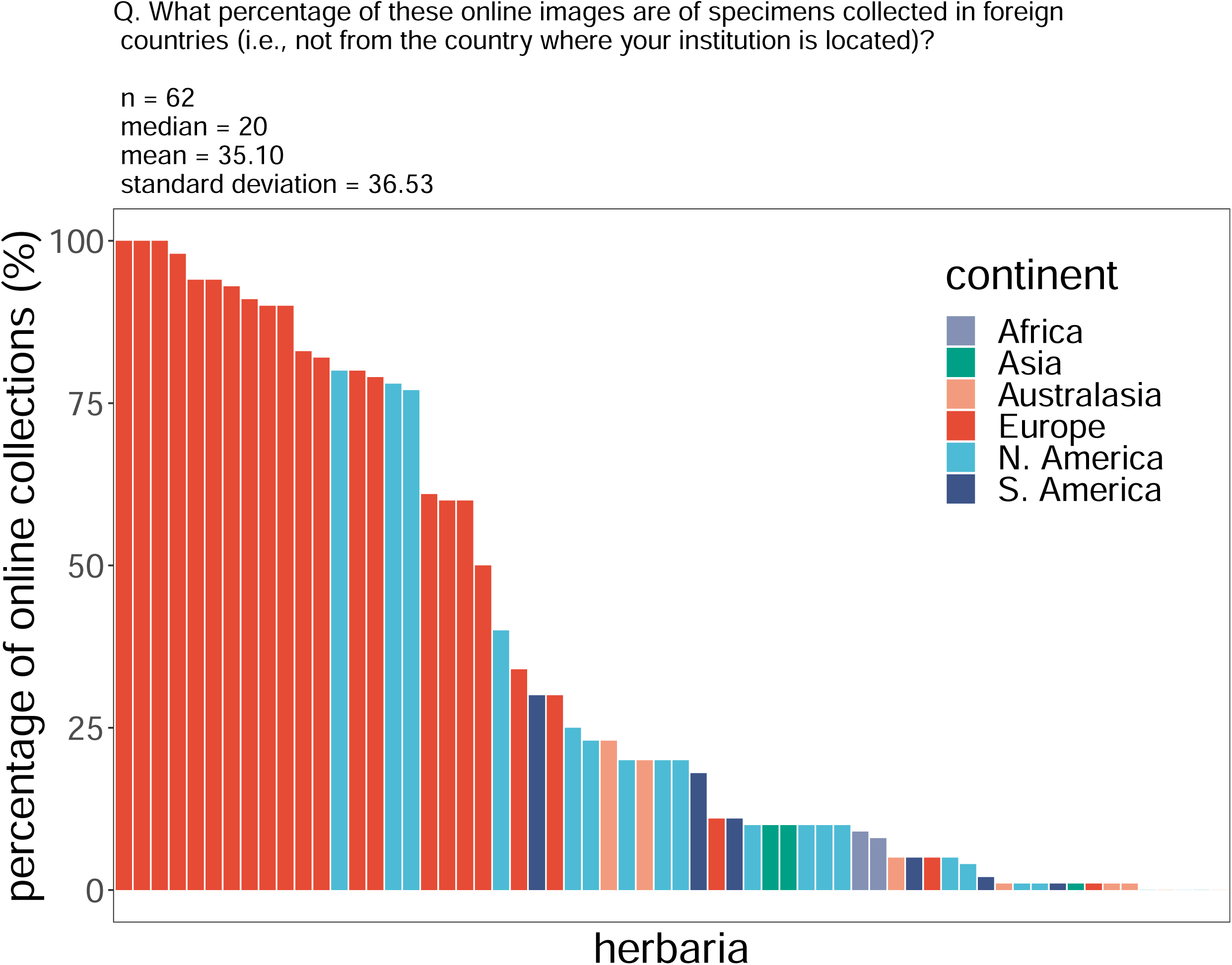

